# A direct fiber approach to model sclera collagen architecture and biomechanics

**DOI:** 10.1101/2022.11.20.517259

**Authors:** Fengting Ji, Manik Bansal, Bingrui Wang, Yi Hua, Mohammad R. Islam, Felix Matuschke, Markus Axer, Ian A. Sigal

## Abstract

Sclera collagen fiber microstructure and mechanical behavior are central to eye physiology and pathology. They are also complex, and are therefore often studied using modeling. Most models of sclera, however, have been built within a conventional continuum framework. In this framework, collagen fibers are incorporated as statistical distributions of fiber characteristics such as the orientation of a family of fibers. The conventional continuum approach, while proven successful for describing the macroscale behavior of the sclera, does not account for the sclera fibers are long, interwoven and interact with one another. Hence, by not considering these potentially crucial characteristics, the conventional approach has only a limited ability to capture and describe sclera structure and mechanics at smaller, fiber-level, scales. Recent advances in the tools for characterizing sclera microarchitecture and mechanics bring to the forefront the need to develop more advanced modeling techniques that can incorporate and take advantage of the newly available highly detailed information. Our goal was to create a new computational modeling approach that can represent the sclera fibrous microstructure more accurately than with the conventional continuum approach, while still capturing its macroscale behavior. In this manuscript we introduce the new modeling approach, that we call direct fiber modeling, in which the collagen architecture is built explicitly by long, continuous, interwoven fibers. The fibers are embedded in a continuum matrix representing the non-fibrous tissue components. We demonstrate the approach by doing direct fiber modeling of a rectangular patch of posterior sclera. The model integrated fiber orientations obtained by polarized light microscopy from coronal and sagittal cryosections of pig and sheep. The fibers were modeled using a Mooney-Rivlin model, and the matrix using a Neo-Hookean model. The fiber parameters were determined by inversely matching experimental equi-biaxial tensile data from the literature. After reconstruction, the direct fiber model orientations agreed well with the microscopy data both in the coronal plane (adjusted R^2^=0.8234) and in the sagittal plane (adjusted R^2^=0.8495) of the sclera. With the estimated fiber properties (C_10_=5746.9 MPa; C_01_=-5002.6MPa, matrix shear modulus 200kPa), the model’s stress-strain curves simultaneously fit the experimental data in radial and circumferential directions (adjusted R^2^’s 0.9971 and 0.9508, respectively). The estimated fiber elastic modulus at 2.16% strain was 5.45GPa, in reasonable agreement with the literature. During stretch, the model exhibited stresses and strains at sub-fiber level, with interactions among individual fibers which are not accounted for by the conventional continuum methods. Our results demonstrate that direct fiber models can simultaneously describe the macroscale mechanics and microarchitecture of the sclera, and therefore that the approach can provide unique insight into tissue behavior questions inaccessible with continuum approaches.

**Highlights:** Collagen fibers are the main load-bearing component of eye tissues.

Conventional sclera modeling ignores that fibers are long, interwoven and interact.

We demonstrate a direct fiber model with long, interwoven and interacting fibers.

Collagen fiber mechanical properties were estimated using inverse fitting.

The model captures simultaneously sclera fiber structure and macroscale mechanics.

## 1. Introduction

Collagen fibers are the principal load-bearing component of sclera (Boote et al., 2020; Girard et al., 2009b; Grytz et al., 2014a; Jan et al., 2017a; Pijanka et al., 2012), and thus play an important role on eye physiology and pathology (Coudrillier et al., 2012; Ethier et al., 2004; Pijanka et al., 2012; Summers Rada et al., 2006). This has motivated many studies aimed at understanding the role of sclera collagen microarchitecture on macroscale eye biomechanical behavior (Coudrillier et al., 2013; Girard et al., 2009a; Grytz et al., 2011).

Because of the complexity of sclera microstructure and the difficulty of accessing it directly for experimentation, numerical models have been widely developed and used for the studies (Coudrillier et al., 2013; Coudrillier et al., 2015b; Girard et al., 2009a; Girard et al., 2009b; Grytz et al., 2011; Hua et al., 2020; Sigal et al., 2004; Voorhees et al., 2018; Voorhees et al., 2017). Most models have been formulated within a continuum mechanics framework in which collagen fiber architecture has been approximated using statistical distributions, often with the parameters derived from inverse fitting. The conventional continuum approach, while helpful to describe the sclera macroscale behavior, does not account for potentially crucial tissue characteristics such as fiber interweaving, fiber-fiber interactions and the in-depth fiber orientation distributions (Boote et al., 2020; Jan et al., 2017b; Lee et al., 2022a). Thus, the conventional approach is limited in the ability to capture sclera structure and mechanics at fiber-level scale. This is problematic because accurate predictions at the small scale are crucial if the intent is to use the models to understand effects at the scale of cells, axons and for studying mechanobiology. Further, ignoring interweaving and fiber-fiber interactions can introduce substantial errors when estimating sclera fiber mechanical properties using inverse fitting (Wang et al., 2020).

To address the limitations of conventional models and better account for microstructure, models have been developed that explicitly incorporate collagen fiber networks (Hadi and Barocas, 2013; Islam and Picu, 2018; Licup et al., 2015; Picu et al., 2018; Zhang et al., 2013). The models, however, are limited in their ability to represent specimen-specific collagen architecture. In addition, the models were formed by short fibers, sometimes generated stochastically, and do not represent well the long fibers that form the sclera (Boote et al., 2020). Long fibers can, potentially, have fundamentally different mechanical behavior than short fibers (Voorhees et al., 2018), and thus that explicitly accounting for them is important in specimen-specific modeling of the eye. Altogether, the limitations of the current modeling tools highlight the need to develop more advanced modeling techniques that can incorporate detailed information on fibers.

Our goal was to demonstrate that it is possible to build a model of sclera that represents fiber microstructure better than the conventional modeling approaches and that also captures sclera macroscale behavior. In this manuscript we introduce a new modeling approach, that we call direct fiber modeling, in which the collagen architecture is accounted for by long, continuous, interwoven fibers. The fibers are embedded in a continuum matrix representing the non-fibrous tissue components. We demonstrate the methodology by modeling a rectangular patch of posterior pole sclera. First, we show that a specimen-specific direct fiber computational model can be built based on high resolution polarized light microscopy data of cryosections from pig and sheep. We show that the fiber orientation distributions of the direct fiber model can be made to follow closely those of the cryosections simultaneously in both the coronal and sagittal planes. This is a more demanding requirement than in conventional models in which only the fiber orientations in coronal plane are accounted for (Girard et al., 2009b). Second, we used the model in an inverse modeling approach by fitting experimental biaxial stress-strain data from the literature. We show that the direct fiber model can match the experiment simultaneously in both radial and circumferential directions. Overall, direct fiber modeling can provide unique insight into the interplay between tissue architecture and behavior and help answer questions that have been inaccessible with the conventional continuum approach.

## 2. Methods

This section is organized in three parts following the same general order as the process for building and using a direct fiber model. First, experimental data on sclera fibers and orientations were obtained using established polarized light microscopy (PLM) imaging of histological cryosections (Jan et al., 2015). Second, a direct fiber finite element model of a patch of posterior sclera was built based on the PLM-derived orientation data. The fiber architecture of the model was first built from the collagen fiber orientation data in the coronal sections, and then iteratively optimized to also match the orientation data from sagittal sections. The direct fiber model was then embedded in a matrix representing non-collagenous components. Third, the combined fiber and matrix model was used in an inverse fitting optimization process to match the models’ simulated stress-strain behaviors with equi-biaxial test data from the literature (Eilaghi et al., 2010). This process produced estimates of the fiber mechanical properties. Below we describe these parts in detail.

Modeling was done in Abaqus 2020X (Dassault Systemes Simulia Corp., Providence, RI, 171 USA). Customized code and the GIBBON toolbox (Moerman, 2018) for MATLAB v2020 (MathWorks, Natick, MA, USA) were used for model pre/post-processing and inverse modeling.

### 2.1 Histology, polarized light microscopy and fiber orientation quantifications

The study was conducted in accordance with the tenets of the Declaration of Helsinki and the Association of Research in Vision and Ophthalmology’s statement for the use of animals in ophthalmic and vision research. For the fiber orientation distribution in the coronal plane, we used a porcine eye that was also part of a previous study on sclera architecture (Gogola et al., 2018b). Details of the eye preparation, histological processing, PLM imaging and post-processing methods are reported elsewhere (Jan et al., 2015; Jan et al., 2017b). Briefly, a healthy eye without known abnormalities was obtained from the local abattoir and processed within 24 hours of death. The episcleral tissues, fat and muscles were carefully removed, and the globe was perfusion and immersion fixed in 10% formalin for 24 hours at an IOP of 0 mmHg. After fixation, the optic nerve head region was isolated with an 11 mm circular trephine and serially cryo-sectioned coronally with a slice thickness of 30 μm (Figure 1 A). Seventeen serial sections were imaged with PLM using the 0.8× objective (NA 0.12) of an Olympus SZX16 microscope, paired with a dual chip Olympus DP80 camera (4.25 μm/pixel).

**Figure 1.**
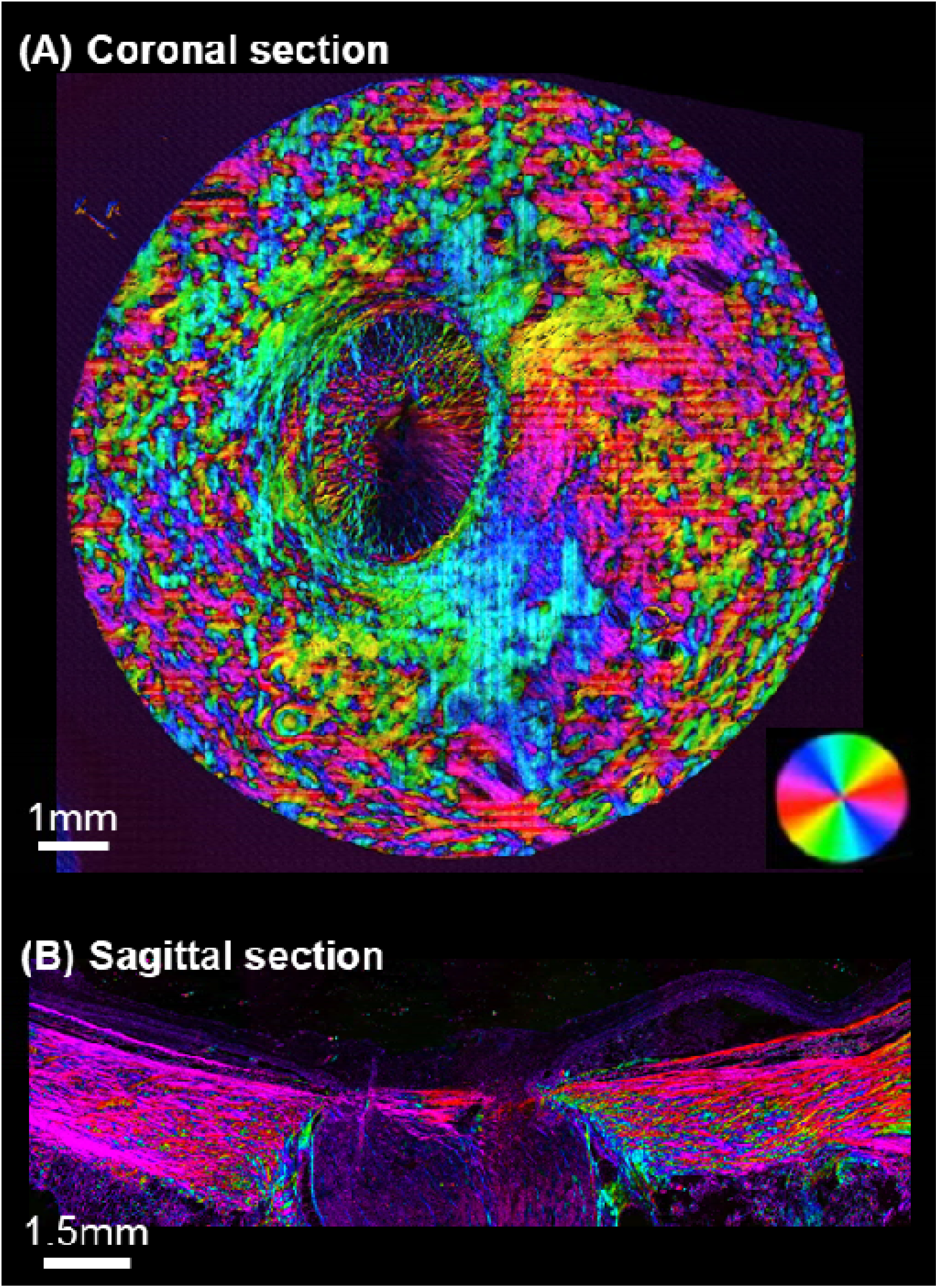
**(A)** Example PLM image of a coronal section of a pig eye through the lamina cribrosa. **(B)** Example PLM image of a sagittal section of a sheep eye through the optic nerve head (ONH). The colors indicate the local fiber orientation in the section plane, and the brightness the “energy” parameter (see main text).

For the fiber orientation distribution in the sagittal plane, we used a healthy sheep eye that was processed in the same way, except that the optic nerve head was sectioned sagittally. Naturally, it is impossible to section the coronal section sample again to obtain the information on in-depth sagittal orientations. Thus, we had to use a different sample. Three sections passing through the posterior temporal sclera and the middle of the scleral canal were selected and imaged with PLM using an Olympus IX83 microscope with 4x objective (1.49μm/pixel). The higher resolution in this orientation was selected to resolve better the in-depth fiber interweaving. Fiber orientation distributions were normalized for use, and therefore we do not expect this to affect the reconstructions. Please see the discussion for a discussion of the potential consequences of having used different species for the coronal and sagittal planes.

PLM images were processed to derive at each pixel the in-plane collagen orientation (in Cartesian coordinates) and a parameter which we have previously referred to as “energy” (Yang et al., 2018b). Energy helps identify regions without collagen, such as outside of the section, and regions where the collagen fibers are primarily aligned out of the section plane, so that they can be accounted for in the orientation distribution.

For the coronal fiber orientation, the coronal sections were stacked sequentially and registered (Gogola et al., 2018b). After registration, the PLM data was reprocessed to align orientation values (Gogola et al., 2018b). As target to build the direct fiber model we selected a rectangular block of sclera, 2.00 x 1.91 mm in size, in the temporal side of the optic nerve head (Figure 2). The location and shape were chosen to match the sample tested experimentally (Eilaghi et al., 2010). We then calculated the distribution of collagen fiber orientations from the PLM images. We used the pixel-level data from 17 sections, weighted by the local “energy”. This allowed us to build the in-plane distribution based on fibers in the same plane (Yang et al., 2018b).

**Figure 2.**
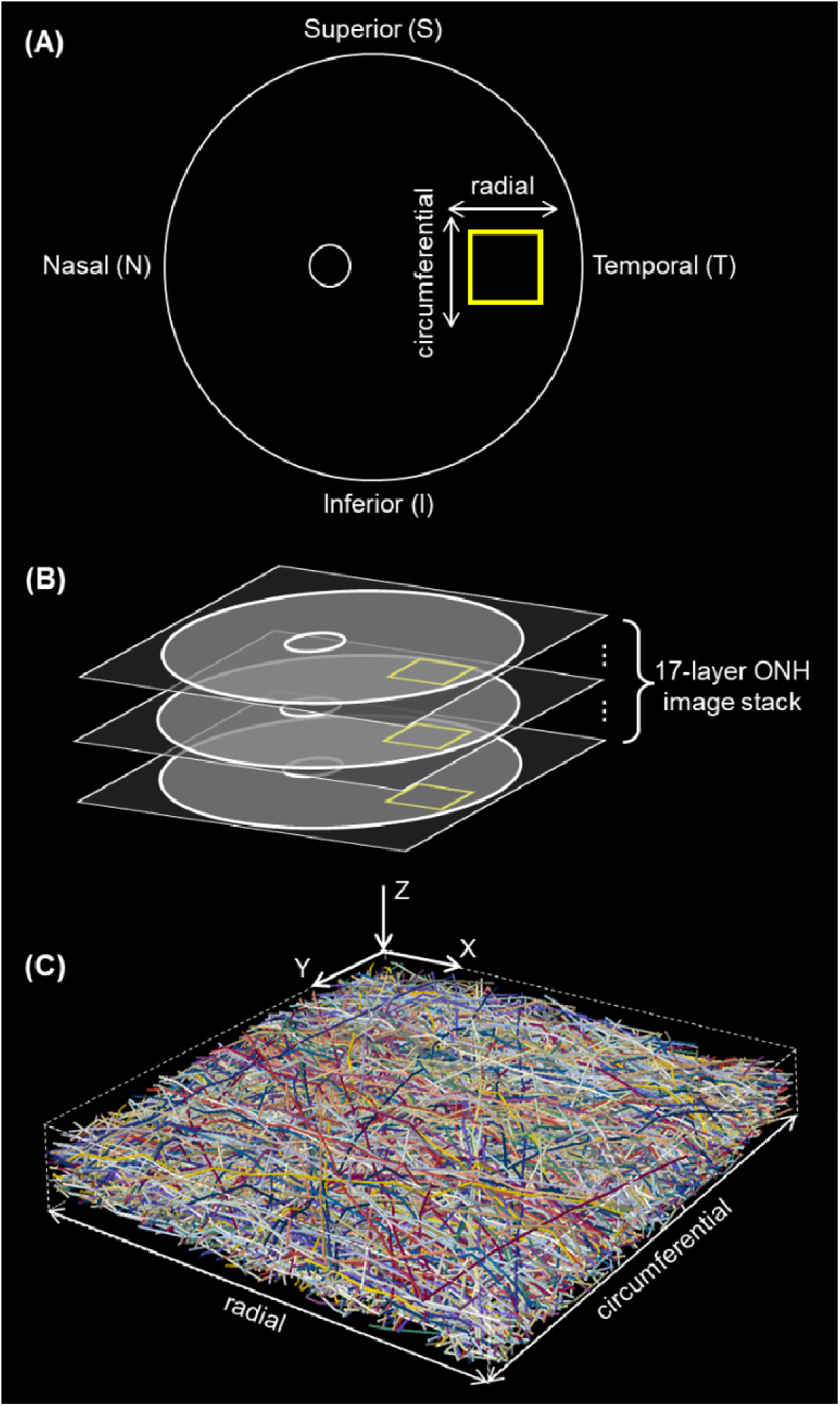
**(A)** Schematic coronal view illustrating the location of the scleral region modeled (yellow box, 2.00 x 1.91 mm). The region was located at the temporal sector. Also shown are the radial and circumferential directions, which correspond with the directions used for the equi- biaxial testing in the experiment and simulation. **(B)** A set of 17 coronal sections were stacked and registered. Fiber orientation data was extracted from the selected rectangular patch of sclera region (yellow box in panels A and B) in the image stack and used to build the direct fiber model **(C)**. Fibers, or fiber bundles, are shown in random colors to simplify discerning the complex interwoven architecture.

**Figure 3.**
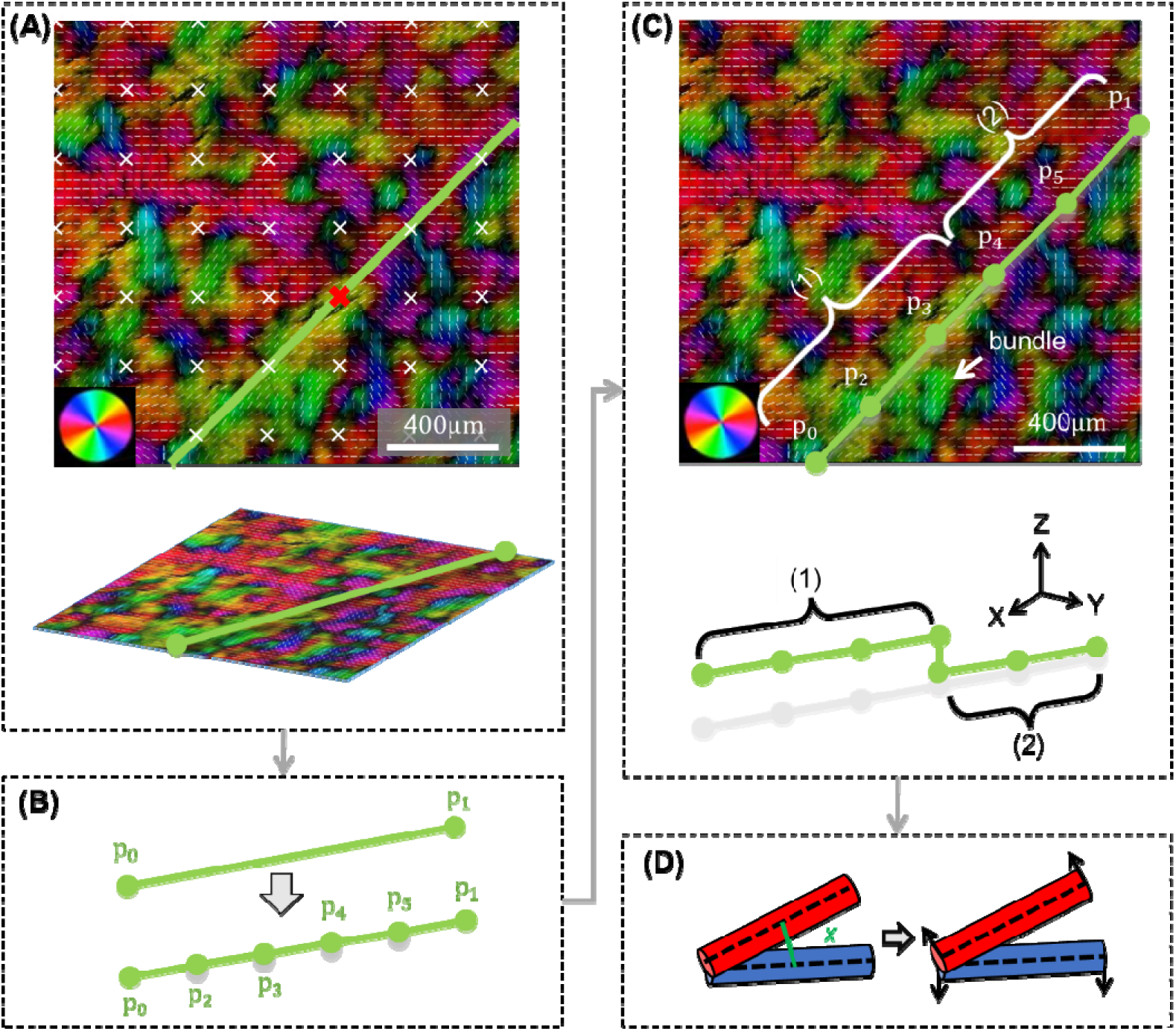
Workflow for creating and processing fibers. **(A)** The process begins with the pixel-level data from the coronal section PLM process (both orientation and energy). For clarity here we illustrate the process with a small square region. The colors of the PLM image represent the local collagen orientation. To help discerning orientations the images are shown with overlaid short white line segments representing the mean orientation over a small square region. (top panel) A regular grid of “seed” points was defined (white x marks). At these seed points the local fiber orientation was sampled, averaged as per the white lines to avoid noise. The energy information was used to skip defining fibers in regions without a reliable well-defined orientation, and to give preference to the in-plane fiber orientations over the out-of-plane ones. The in-plane orientation was then used to define a straight fiber in the image plane. In the example case, a fiber was traced at the grid point (red x mark) at an angle around 45 degrees. The bottom panel shows an isometric view of the image with the fiber overlaid. **(B)** (top) The fiber was initially defined by one element and two nodes, p_0_ and p_1_. (bottom) The fiber was then refined into five elements and six nodes. **(C)** The average orientations over the fiber elements were then computed and compared with local image orientations to determine their agreement element by element. In the example shown, the element group (1), p_0_-p_2_, p_2_-p_3_, p_3_-p_4_, agreed well with the direction of a yellow-green bundle. Accordingly, the group was accepted at this section. In contrast, the element group (2) p_4_-p_5_, p_5_-p_1_, had poor agreement between element and image orientations. This group was then assigned a lower depth, effectively “pushing” the group or fiber segment to the depth of another section. Fiber connectivity was ensured by adding an element to connect the two element groups ((1) and (2)) at different depths. The fiber smoothness was restored when resolving fiber collisions. Meanwhile, elements were combined or split to maintain all element lengths within a pre-defined range. **(D)** Elements were moved apart if the shortest distance x between them was smaller than fiber diameter, indicating a collision.

For the sagittal fiber orientations distributions, we sampled sections through the posterior temporal sclera where the coronal information was obtained. However, instead of selecting one region and section for sampling, we reasoned that it was better to obtain a sagittal fiber orientation distribution that was more broadly representative of the region. This would help ensure good correspondence with the orientation distribution in the coronal orientation, and it would account for potential variations between eyes and misalignment between regions. To obtain the representative distribution, we measured the orientation distribution in 236 regions of three sagittal sections. All of these measurements were in regions 715 μm x 715 μm in size, and close to the location from where the coronal sample was obtained. In each of those regions, the orientation distributions were calculated by using the out-of-plane information derived from the energy parameter to restrict the measurements to be on sagittal plane (Yang et al., 2018b). We then computed the average orientation distribution over all these. We also calculated the sagittal fiber anisotropy from the median of the anisotropies among the 236 areas. The anisotropy indicates the degree of fiber alignment and was quantified as circular standard deviation (Gogola et al., 2018b). Perfectly aligned fibers have an anisotropy of 1 and evenly dispersed fibers have an anisotropy of 0. We reasoned that the average distribution and median anisotropy provide a better representation of the orientation distribution in this direction than any single distribution.

### 2.2 Direct fiber model construction

#### 2.2.1 Fibers

Fibers were simulated using 3-dimensional linear truss elements (T3D2 in Abaqus). Fiber locations were defined by Cartesian coordinates (X, Y, Z) of element nodes. To define fibers, we sampled orientation values from PLM images at regularly spaced “seed” points with a spacing of 272 μm. At each seed point, a straight fiber 8.5 μm in diameter was traced in the section plane at an angle equal to the orientation at the seed point, meanwhile, the fiber passed through the seed point and extended the full span of the region. The process was applied for each layer, and then the fibers were stacked, resulting in a stack of 2D layers of fibers, each with a large number of fibers crossing or “interpenetrating”. To resolve fiber interpenetrations, an algorithm was used to refine and displace fiber elements in the whole structure (Matuschke et al., 2021; Matuschke et al., 2019). Briefly, if the smallest distance between two elements was less than the fiber diameter, an interpenetration was detected. Interpenetrated elements were shifted apart iteratively until all interpenetrations were resolved. During the process, fiber elements were re-meshed so that the element length was kept between 25.5μm to 51μm. Fibers were smoothed by controlling the fiber minimum radius of curvature.

It is important to make a few important notes regarding our use of the term “fiber”. The collagen of the sclera has a complex hierarchical structure even more complex than that of the cornea and tendon (Boote et al., 2020; Jan et al., 2018). The models were reconstructed using a fiber diameter of 8.5 μm, which means that what we refer to as a fiber most likely represents a group of fibers that elsewhere may instead be described as a fiber bundle. Nevertheless, because our intent with this work is to call attention to the power of incorporating detailed microstructural information on sclera, we decided to use the term fiber, while also being careful to acknowledge at several critical points that these may be understood as fiber bundles. This is further addressed in the discussion.

To evaluate the similarity of the model and the images in the coronal plane, we quantified the model’s coronal orientation distribution and compared it with the distribution of the PLM images. We counted the occurrences of element orientations, where the orientation of an element was the slope angle in the coronal section plane and the number of occurrences was estimated as the element volume. This allowed us to account for uneven element sizes and to properly compare with PLM pixel-based measurements.

To better account for the fiber distribution in 3D, including the through-depth fiber undulations, we did the following: we shifted the fiber elements locally in-depth according to the fiber relative positions observed in coronal section images. If a fiber element had an orientation that disagreed with the orientation in the image by more than 45 degrees, the element was “shifted” in the posterior direction. A fiber segment that was in agreement with the local orientation was kept at the original sclera depth. After applying this process throughout the volume, we did another iteration resolving fiber interpenetrations. We then compared the in-depth fiber anisotropy of the model with the representative anisotropy in the sagittal plane. If the model had a larger in-depth anisotropy than the images, we shifted the fibers, increasing the fiber undulation amplitudes and decreasing the degree of fiber alignment. Conversely, if the model had a lower anisotropy than the sagittal images, we decreased the amplitude of fiber undulations. Each of these steps was followed by an iteration resolving fiber interpenetrations and a check in the agreement between the model and sample anisotropies in the coronal plane. Adjusted R-square (adjusted R^2^) values were used to evaluate the fitness of orientation distributions. This process converged to a set of continuous fibers in excellent agreement with the PLM-derived orientations simultaneously in both coronal and sagittal planes. Adjusted R^2^ is similar to the conventional R^2^ in that it indicates fit to a curve, but it adjusts for the number of points considered, avoiding potentially misleading excellent fits that are the result of many points of comparison (Miles, 2005).

#### 2.2.2 Matrix

The model used for inverse fitting consisted of the direct fiber model and a coincident matrix with an overall dimension of 2.00 x 1.91 x 0.35 mm. The model of fibers and matrix had the same dimensions, and fiber end-nodes resided on the matrix surfaces.

### 2.3 Model inverse fitting

#### 2.3.1 Meshing and material properties

Fibers were modeled as a hyperelastic Mooney-Rivlin material (Holzapfel, 2001):

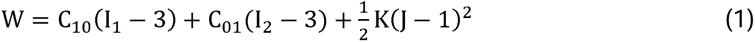

where W was the strain energy density, C_10_ and C_01_ were the material constants, restricted by C_10_ + C_01_ > 0 and would be determined by inverse modeling, I_1_ and I_2_ were the first and second invariants of the right Cauchy-Green deformation tension, K was the bulk modulus and J was the determinant of the deformation gradient.

Fibers were simulated using 3-dimensional linear truss elements (T3D2 in Abaqus). A mesh convergence analysis was performed to verify that both the fiber and matrix models were sufficiently refined. The results shown were obtained with fiber element lengths between 25.5μm to 51μm, resulting in element aspect ratios from 3 to 6. This is consistent with a previous study on fiber mechanics in which we found that aspect ratios of 5 were optimal (Islam and Picu, 2018). The matrix was meshed with linear eight-nodded, hybrid hexahedral elements (C3D8H in Abaqus) and modeled as a neo-Hookean material with a shear modulus of 200 kPa (Coudrillier et al., 2015a; Girard et al., 2009b). The element size of the matrix was 0.0887mm.

#### 2.3.2 Interactions

Fiber-fiber interactions were simulated by preventing fiber interpenetrations using Abaqus’ general contact with no friction. Preventing fiber interpenetration is computationally expensive and therefore commercial implementations are often directed at maintaining small interpenetrations rather than avoiding them altogether. We reasoned that a small interpenetration, in low single digits percent of fiber diameter, will likely not represent a major deviation from fiber physical behavior. We calculated the extent of the interpenetration in the following way: we defined a deformed model based on the original model and the final nodal displacements. A customized MATLAB program was then used to find all existing interpenetrated element pairs. For each of these pairs, we quantified the percentage of interpenetration as:

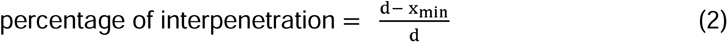

where d was fiber diameter, and x_min_ was the minimum distance between two elements.

Interpenetration thus varied between 0 when there is no interpenetration and 1 when two elements are perfectly overlapping. We found that 99.4% of the interpenetrations were smaller than 5%. This is further addressed in the discussion.

Fiber-matrix interactions were ignored, as is usual in biomechanical models of the eyes (Grytz et al., 2014a; Grytz and Meschke, 2009; Petsche and Pinsky, 2013).

#### 2.3.3 Finite element analysis procedure

The fiber-matrix assembly was subjected to a quasi-static process of equi-biaxial stretch to match the experimental results and boundary conditions reported elsewhere (Eilaghi et al., 2010). The matrix was simulated using Abaqus standard implicit procedure. Due to the presence of complicated fiber contacts, the direct fiber model was simulated using Abaqus dynamic explicit procedure, meant to improve the convergence and computational efficiency. The model fiber volume fraction (VF) is 7%. The resulting stresses σ along radial and circumferential directions were contributed by both matrix and fibers, where the matrix stress was weighted by fiber VF:

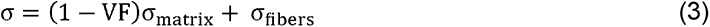

Because the process was quasi-static, the dynamic explicit procedure would require an excessive number of small-time increments, which is computationally impractical. Therefore, to run the dynamic analysis efficiently, mass scaling was implemented. We achieved a 1e-5 stable time increment, where we modeled the process in the shortest time period in which inertial forces remain insignificant. During the simulation, we assured that the inertial effects were negligible by keeping the kinetic energy less than 5% of the internal energy.

#### 2.3.4 Boundary conditions and inverse modeling procedure

The model was simulated iteratively to derive fiber material properties (C_10_ and C_01_) by inversely matching published experimental stress-strain data (Eilaghi et al., 2010).

##### Selecting the target experimental set

The fibers in the direct fiber model were not equally distributed in all directions. It seems reasonable to expect that this structural anisotropy will result in mechanical anisotropy (Coudrillier et al., 2015b). It was therefore important to select for inverse fitting, among the multiple experimental results reported in the literature (Eilaghi et al., 2010), the one that exhibited a mechanical anisotropy matching the structural anisotropy of our model. To do this, we first quantified the mechanical anisotropy of our model (Figure 4). We did this by subjecting the model to displacement-controlled equi-biaxial stretch tests. At each strain level we quantified the model’s stress anisotropy as the ratio of stresses in the orthogonal directions (S_11_/S_22_). The test was repeated several times with different fiber material property values. The results showed that the ratio of stresses had a fairly constant distribution and was essentially independent to the material properties changes within the ranges tested. With this, we were able to select the case with anisotropy most similar to that of our model among the several experimental results reported by Eilaghi et al.. Specifically, we selected the data for sample 76940-TS, as labeled by Eilaghi et al.. (Eilaghi et al., 2010)

**Figure 4.**
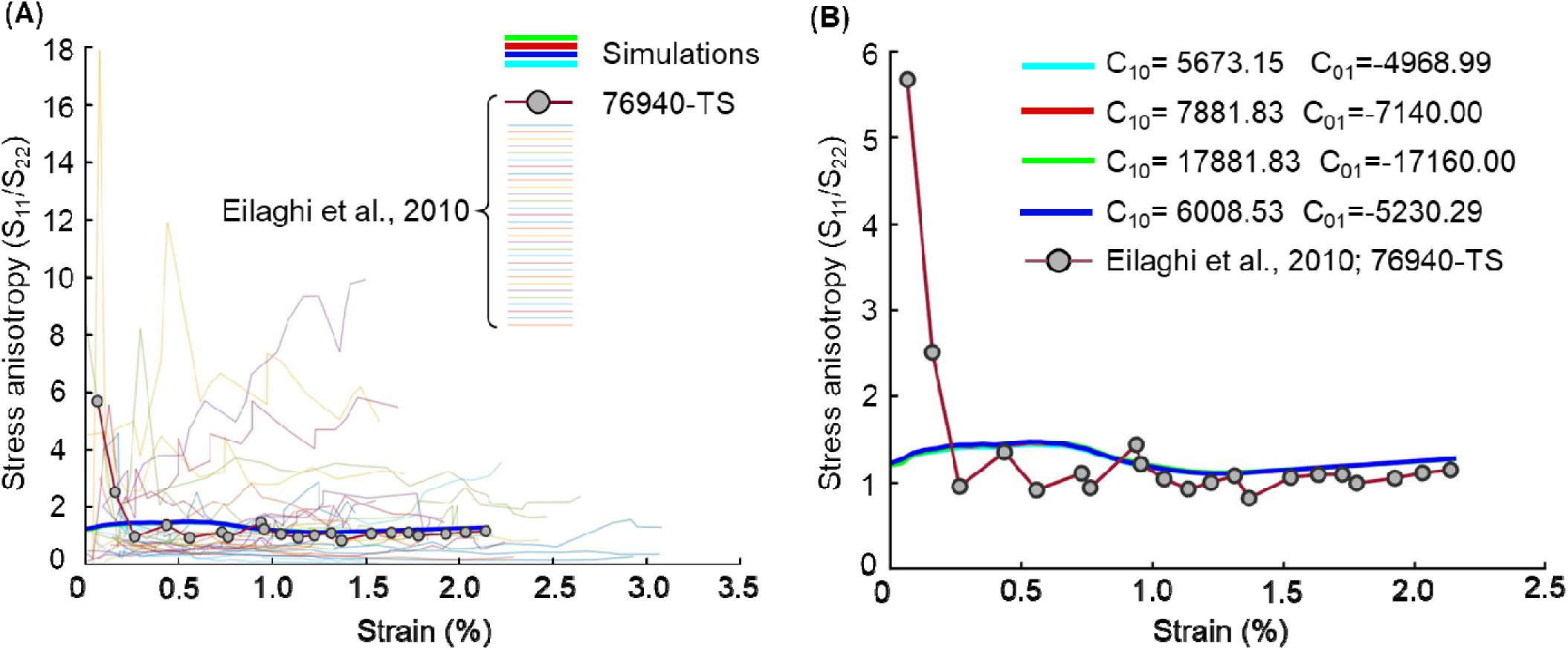
**(A)** To properly fit the model to an experimental curve it was necessary to select out of all the experimental curves the set with a similar anisotropy. See text for more details. To do this we first assigned four sets of random material parameters, C_10_ and C_01_, to the fiber model. The intent was to generate a set of model behaviors spanning the range of anisotropies to be expected over a wide range of material properties. Equi-biaxial stretch was applied to the fiber-matrix assembly and the stress anisotropy (S_11_/S_22_) of the model computed at many steps through the stretching process. We then also computed the stress anisotropy of the experimental curves reported by Eilaghi et al. There were substantial variations in their anisotropies. Out of all of the curves we selected 76940-TS because it had the closest agreement with the model anisotropy. The plot shows all experimental curves with 76940-TS highlighted. There was good agreement in anisotropy after about 0.25% strain. At smaller strains the experiment observed higher anisotropy, potentially due to the challenge of balancing the initial loads in the clamping when using rakes. **(B)** In this plot only the four models and the selected experimental curve. All C_10_ and C_01_ values are in MPa. The results revealed that the model stress anisotropy was essentially independent of the fiber material properties. This can be discerned from the observation that the lines representing the four simulations are almost indistinguishable. This means that the process of fitting fiber material properties will preserve the stress anisotropy, allowing a close match of the stress-strain data simultaneously in both radial and circumferential directions.

To be consistent with the selected experimental data, we assigned an equi-biaxial stretch of 2.16% as the displacement boundary condition to our fiber-matrix assembly. To optimize the fiber material properties (C_10_ and C_01_), we used the simplex search method of Lagarias et al. (Lagarias et al., 1998). The algorithm sought to identify the two parameters that yielded the closest match between the simulated and experimental stress-strain curves, simultaneously in both directions. We optimized the fitness by minimizing the cost function, defined as the residual sum of squared (RSS) between the model and experimental curves at 21 strain states. The optimization was terminated when the cost function value was smaller than 0.01. After optimization we also computed the adjusted R^2^ between the curves to assess curve similarity. The complete inverse fitting process was repeated 11 times with various starting parameters to test the consistency and “uniqueness” of the results.

For interpretation of the results and to compare with other studies, it is useful to “convert” the optimal C_10_ and C_01_ parameters of the hyperelastic Mooney-Rivlin model into more intuitive representations of fiber mechanical properties. Specifically, we estimated the fiber shear modulus as 2(C_10_ + C_01_). We also derived a fiber elastic modulus by simulating uniaxial stretch of a single straight fiber with the optimal C_10_ and C_01_ values and the hyperelastic Mooney-Rivlin material. The fiber elastic modulus was then obtained as the slope of the stress-strain curve at 2.16% strain.

## 3. Results

After construction, the model consisted of 1016 fibers (or fiber bundles). Fiber orientation distributions and overall anisotropies were similar between the model and the tissue, as measured using PLM (Figure 5). In the sagittal plane, the anisotropies were 0.6031 and 0.5976 for the model and tissue, respectively (a difference smaller than 1%); In the coronal plane, the anisotropies were 0.1777 and 0.1626 for the model and tissue, respectively (a difference smaller than 10%).

**Figure 5.**
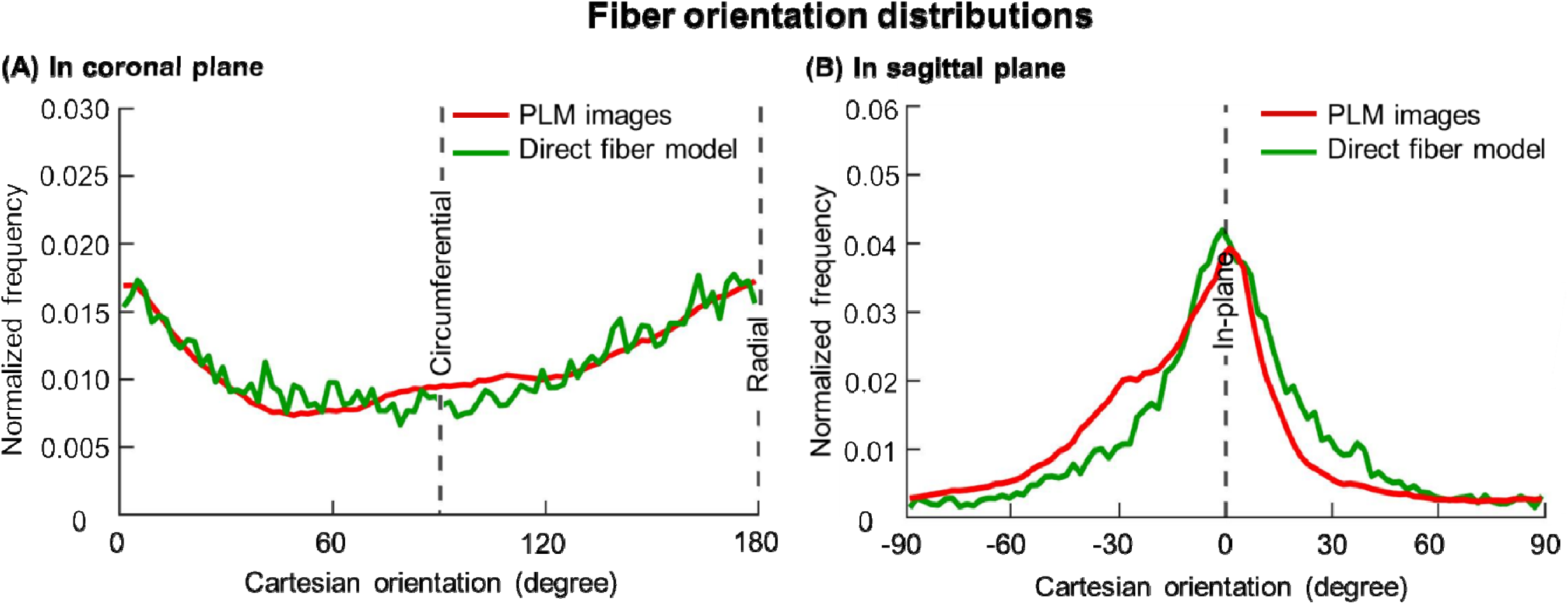
Fiber orientation distributions of the direct fiber model (green lines) and the PLM images (red lines), in the **(A)** coronal and **(B)** sagittal planes. For the coronal plane the PLM orientation was derived from the stack of 17 images. In the coronal plane the radial direction corresponds to 0 and 180 degrees and the circumferential direction with 90 degrees. For the sagittal plane, the PLM orientation shown is the average orientation distribution over the 236 regions analyzed. See main text for details. Frequencies were normalized by the total sum of frequencies. Overall these results show that fiber orientation distributions of the direct fiber model agreed well with those from the PLM images in both coronal (adjusted R^2^ = 0.8234) and sagittal (adjusted R^2^ = 0.8495) planes.

Figure 5 shows the fiber orientation distributions of the direct fiber model and PLM images. The match of distributions was achieved simultaneously in both coronal and sagittal planes. By Wilcoxon rank sum tests, there were no significant differences in fiber orientation distributions between the model and the PLM images, in either coronal (p > 0.7) or sagittal (p > 0.6) planes.

Figure 6 illustrates fiber displacements and stresses at half and full stretch. The large differences in stress between fibers were consistent with the anticipated process of stretch-induced recruitment (Grytz and Meschke, 2010; Jan and Sigal, 2018). All 11 optimization runs led to curves that were in good agreement with the experiment. They all led to fairly consistent estimates of fiber shear modulus, with an average of 1522.8 MPa and a standard deviation (STDEV) of 38.9 MPa (Supplementary Table 1). Interestingly, these were obtained with C_10_ and C_01_ coefficients that varied substantially. For example, C_10_ ranged from 118.9 to 7881.8 MPa.

**Figure 6.**
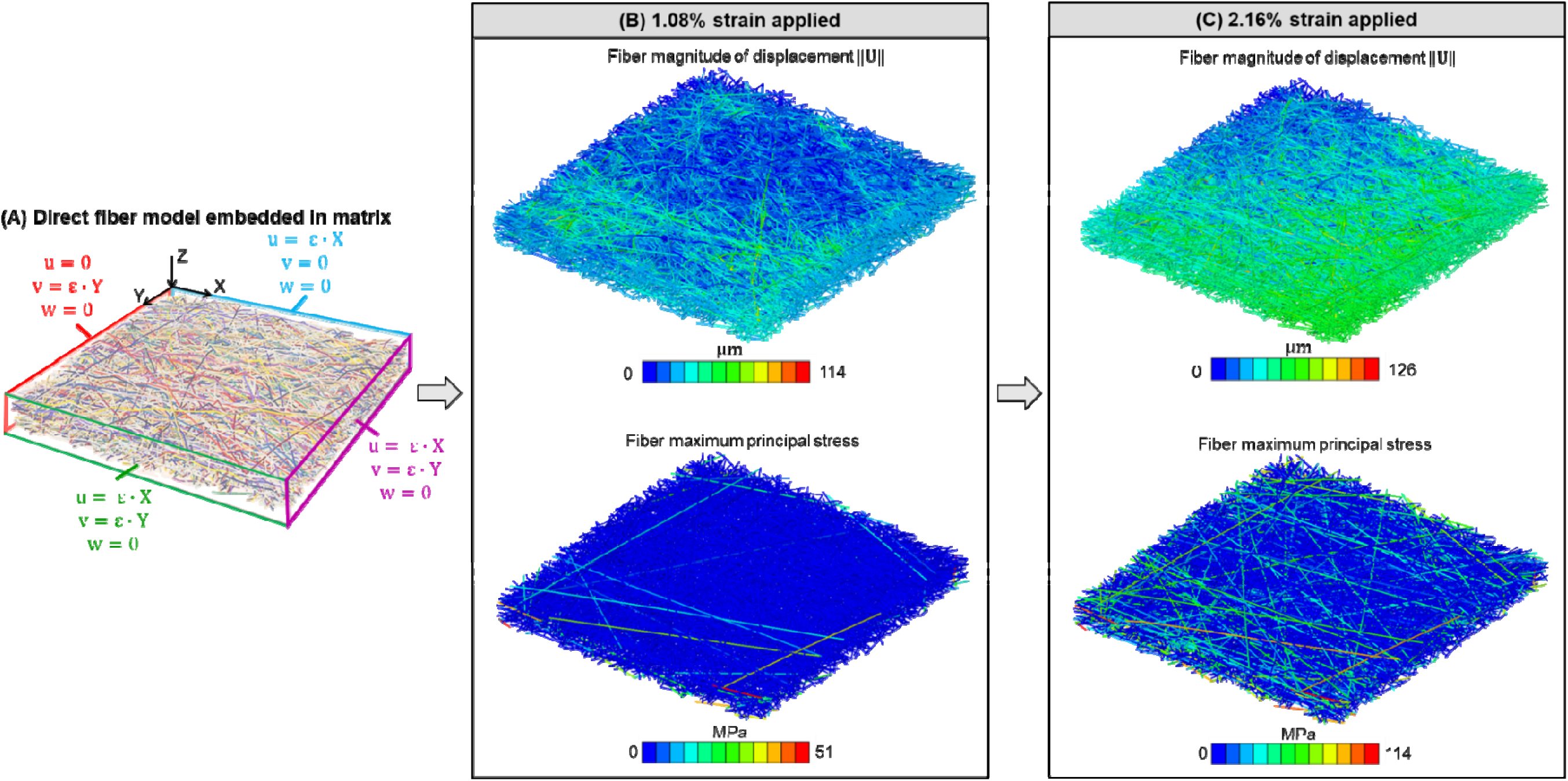
**(A)** Schematic of the displacement boundary conditions for the fibers and matrix to represent scleral tissue subjected to 2.16% equi-biaxial strain (). **U** is the nodal displacement vector of fiber with components u, v & w representing displacement in X, Y and Z direction, respectively. is the fiber magnitude of displacement. In panel A the fibers are randomly colored to facilitate discerning their architecture. Panels **B** and **C** show isometric views of the direct fiber model with the fibers colored according to the magnitudes of displacement (top row) or maximum principal stress (bottom row). The model is shown when subjected to **(B)** 1.08% strain or **(C)** 2.16% strain. From the images it is clear that the model fibers exhibit complex loading patterns.

Figure 7 shows that the stress-strain curves of the optimal model fit very well the experimental data in both radial (adjusted R^2^ = 0.9971; RSS = 0.00095) and circumferential directions (adjusted R^2^ = 0.9508; RSS = 0.0098) simultaneously. The material parameters for the inverse fitting run that led to the closet model and experiment fit were:

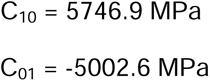

**Figure 7.**
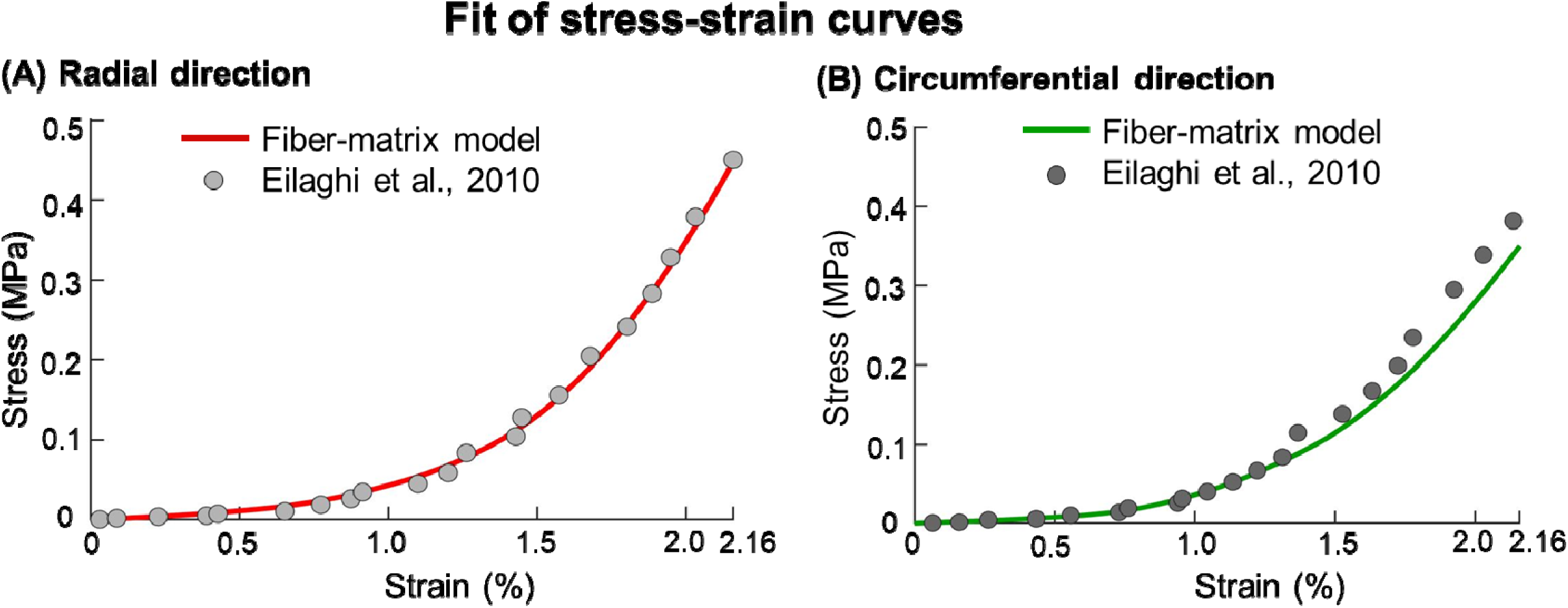
Fit of stress-strain response was achieved between the model and the experiment simultaneously in both the (A) radial direction (adjusted R^2^ = 0.9971; RSS = 0.00095) and (B) circumferential directions (adjusted R^2^ = 0.9508; RSS = 0.0098)

The estimated fiber elastic modulus at 2.16% strain was 5.45GPa. The estimated fiber shear modulus is 1488.6 MPa.

## 4. Discussion

Our goal was to develop a direct fiber modeling approach that captures both the microscale collagen fiber architecture and the macroscale mechanical behavior of the sclera. We demonstrated our approach by modeling a rectangular patch of the posterior sclera. The model incorporated several fiber characteristics ignored by most previous models of posterior sclera, such as fiber interweaving, fiber-fiber interactions, long fibers, and a physiologic experimentally-derived in-depth fiber orientation distribution. Below we provide a brief overview of the use of conventional continuum approaches to model the sclera, followed by a discussion on the importance of considering the fiber characteristics described above, and the strengths of the direct fiber modeling approach. We continue with a discussion on how to further interpret our models and the results obtained from them, including a comprehensive list of limitations.

### Conventional continuum models of sclera

Numerical models of sclera have been widely developed and advanced for studying sclera structure and mechanics (Coudrillier et al., 2013; Coudrillier et al., 2015b; Girard et al., 2009a; Girard et al., 2009b; Grytz et al., 2011; Hua et al., 2020; Sigal et al., 2004; Voorhees et al., 2018; Voorhees et al., 2017). Most models have been formulated within a continuum mechanics framework in which collagen fiber architecture has been approximated using statistical distributions. For example, in a common approach, the scleral collagen microarchitecture is accounted for by collagen fiber “families”, each of which is described through the family preferred orientation and a “degree of alignment” around this preferred orientation (Coudrillier et al., 2013; Girard et al., 2009a; Grytz et al., 2011). More advanced models have incorporated experimental data on collagen fiber orientations, obtained for example from small-angle light scattering or related scattering techniques (Coudrillier et al., 2013; Schwaner et al., 2020a; Zhang et al., 2015). Other recent models have focused on incorporating regional variations in fiber family characteristics (Kollech et al., 2019). The success of the conventional models at the macroscale encouraged their use to predict microstructural characteristics of sclera fibers, such as collagen fiber crimp (Girard et al., 2009a; Grytz et al., 2014a; Grytz et al., 2011).

There is no doubt that the conventional models have been helpful in capturing macro-scale sclera mechanics, and to understand the interplay between mechanics and structure, both micro and macro. However as the experimental tools advance, the information available on sclera microstructure and mechanics has improved (Behkam et al., 2019; Brown et al., 2007; Coudrillier et al., 2016; Hoerig et al., 2022; Jan et al., 2015; Lee et al., 2022a; Ling et al., 2019; Pijanka et al., 2019; Sigal et al., 2014; Winkler et al., 2010; Yang et al., 2018a; Yang et al., 2021). These studies have demonstrated structural and mechanical characteristics of the sclera that are potentially crucial yet are not accounted for by the conventional continuum approach for modeling, such as fiber interweaving, fiber-fiber interactions and the in-depth fiber orientation distributions (Boote et al., 2020; Jan et al., 2017b; Lee et al., 2022a). More importantly, some of those fiber characteristics are not suited to be incorporated under the continuum framework. This brings to the forefront the need to develop more advanced modeling techniques that can incorporate and take advantage of the newly available highly detailed information.

### The importance of considering fiber interweaving/fiber-fiber interactions, long fibers, and in-depth fiber orientation distribution

A common preconception about the effects of fiber interweaving, and the resulting fiber-fiber interactions, seems to be that interweaving increases the stiffness of a material, or tissue. However, by comparing models with interweaving and non-interweaving fibers, we have shown that an interwoven architecture is, as a structure, more compliant than a non-interwoven architecture (Wang et al., 2020). This is consistent with the literature on textile mechanics (Saiman et al., 2014; Stig and Hallström, 2019). This effect can be understood by noting that the fibers of the interweaving architecture are undulated, whereas those from the non-interweaving models are straight. Straight fibers aligned with the load carry the forces more efficiently, and are shorter, and thus, the overall model is stiffer. Since fiber interweaving and the resulting fiber-fiber interactions play an important role in determining the structural stiffness of the sclera, it should be considered when modeling sclera biomechanics, as did our direct fiber model. Note that we are not the first to recognize the importance of fiber interweaving and fiber-fiber interactions in soft tissue biomechanics (Elliott and Setton, 2001; Guerin and Elliott, 2007; Nerurkar et al., 2011; Wagner and Lotz, 2004; Zhang et al., 2013). For example, in annulus fibrosus of intervertebral disc, interlamellar shearing can account for nearly 50% of the total stress associated with uniaxial extension. Therefore, interweaving collagen fiber layers may play an important role in annulus fibrosus tissue function.

It is important to note that the scale of the undulations of interweaving is different from that of the collagen fiber waviness referred to as crimp by us, (Gogola et al., 2018a; Jan et al., 2018; Jan et al., 2017a; Jan and Sigal, 2018) and others (Grytz et al., 2014a; Grytz et al., 2020a; Grytz and Meschke, 2009). For example, ocular collagen fibers crimp has a period typically under 20 μm, (Gogola et al., 2018a; Jan et al., 2018; Jan et al., 2017a; Jan and Sigal, 2018) much smaller than the estimates of interweaving undulations on the order of 100 to 300 μm (Wang et al., 2020). From a mechanical perspective, the undulations of crimp affect directly the biomechanical behavior of a fiber under load, with the crimp “disappearing” as the fiber is loaded or stretched, and eventually recruited (Jan et al., 2022; Jan and Sigal, 2018). The interactions between interwoven fibers mean that these undulations do not “disappear” under load, and are limited by the interlocking (Lee et al., 2022b).

The collagen fibers of the sclera are long and continuous; thus, they can transfer forces over a long distance (Boote et al., 2020; Voorhees et al., 2018). However, the conventional continuum approach to modeling sclera only considers the local or regional orientation distribution of these long fibers and maps such information into fairly small “finite elements” (Coudrillier et al., 2013; Grytz et al., 2011). As a result, the force transmission is continuous across elements rather than along fibers, which alters the predicted mechanical behavior locally and potentially at the macroscale (Campbell et al., 2015; Coudrillier et al., 2013; Kollech et al., 2019; Roberts et al., 2010; Voorhees et al., 2018; Zhang et al., 2015; Zhou et al., 2019). We are not the first to recognize the importance of accounting for long fiber mechanics (Grytz et al., 2020b; Huang et al., 2017; Lanir, 2017). One approach proposed to address the problem has been through the use of symmetry boundaries across elements. In this manner it is possible to simulate fibers that are continuous across elements. The approach, however, substantially limits the type of models that can be considered compared with the direct fiber model we present.

The collagen architecture of the sclera varies in depth (i.e., the direction perpendicular to the scleral surface) (Jan et al., 2017b; Pijanka et al., 2015). Such variations are crucial in the load-bearing capacity of the sclera (i.e., bearing shear stresses), and may even have clinical implications (Danford et al., 2013). Unfortunately, most fiber-aware models of the sclera only account carefully for the fiber orientations within the scleral plane, while the in-depth orientations are ignored or are modeled in a much more simplified manner than the out of plane orientations (Coudrillier et al., 2013; Voorhees et al., 2017; Zhang et al., 2015). Our direct fiber model incorporates both in-plane and in-depth specimen-specific fiber orientation distributions that match those measured using PLM, and thus, has a higher fidelity for representing the sclera and its mechanics.

### Strengths of direct fiber modeling

Ultimately, a model is only as good as the predictions that can be made using it. On this, we point out that the fiber elastic modulus estimated with our direct fiber model is well within the range of values reported in the literature. For example, we estimated a fiber elastic modulus of 5.45 GPa, compared with those of 2-7 GPa of the bovine Achilles tendon (Van Der Rijt et al., 2006; Yang et al., 2007) and 5-11.5 GPa of the rat tail (Wenger et al., 2007). In contrast, the estimates of fiber elastic modulus derived using continuum models, which used constitutive model formulations based on highly simplified assumptions of fiber architecture and behavior at the microscale level, are several orders of magnitude smaller, between 1 and 200 MPa (Coudrillier et al., 2015b; Grytz et al., 2014b; Schwaner et al., 2020a; Schwaner et al., 2020b). Hence, whilst both continuum and direct fiber models can closely approximate the macroscale sclera behavior, the estimates of fiber mechanical properties derived from both types of models can be substantially different. This may not be a problem when the intent is limited to describing tissue mechanical response. However, an important application of continuum models is to “infer” characteristics of the underlying fibers. The large differences in fiber properties estimated by the continuum models and the experimental measurements already suggest that these should be interpreted very carefully as the values inferred may be inaccurate. A common application of this approach is to compare the fiber properties inferred from healthy and unhealthy tissues, or from young and old donors. The argument in this case is based on the idea that the methods may not produce accurate fiber estimates, but that the comparison remains valid. This may indeed be the case. But it is also possible that if the changes with pathology or age involve aspects of the fibers that are not accounted for in the continuum model formulation, then the origin of the changes will end up artefactually attributed to another tissue characteristic. Thus, while all models involve important approximations, if the goal is to derive estimates at the fiber scale, we posit that direct fiber models are preferrable over continuum models.

The methodology to build the direct fiber model structure can be applied to other collagenous tissues. The reconstruction method is not tied to PLM and can instead be done using second harmonic imaging or confocal microscopy, as long as they provide information on the density and orientation of collagen fibers. The level of detail necessary from the images can depend on the complexity of the tissue in question and the accuracy needed from the reconstruction and simulations. Because the sclera has a complex 3D architecture, we combined histological information from two sectioning directions. This may not be necessary for other tissues, or perhaps it may be possible to obtain depth information by taking advantage of the confocal nature of the imaging or 3D PLM (Yang et al., 2018b).

### Considerations of the uniqueness of material parameters and fiber interpenetrations

A common concern with inverse fitting is the uniqueness of the so-called optimal parameters (Girard et al., 2009a; Girard et al., 2009c; Zhou et al., 2019). When we repeated the optimization with various starting parameters, we obtained fairly consistent fiber shear modulus predictions. However, the material model parameters C_10_ and C_01_ varied substantially. This was not a surprise given the material formulation we used and the clear potential for interactions between the parameters (van Tonder et al., 2023). Altogether, this reinforces the importance of focusing on parameters with a clear physical interpretation, in this case fiber shear modulus. Such parameters are more likely to have better stability.

From a numerical perspective, prevention of fiber penetrations is computationally intensive. We took advantage of highly mature general contact tools implemented in commercial code to keep solution time reasonable. Ensuring absolutely no interpenetration would have required an impractical number of small elements to represent the undulating fibers. To avoid this problem, we decided to quantify and track the interpenetrations and consider a solution valid if these remained below 5% of the fiber diameter. In a worst-case scenario this would mean that two interacting fibers have “pushed” into each other such that the distance between their centers is only 95% of twice their radii. We assumed that the fibers are all circular and remain circular, despite pushing into each other.

## Limitations

When interpreting the results from this work, it is important to consider also the following limitations. First, we considered highly simplified fiber-fiber interactions and ignored fiber-matrix interactions. We are aware that the forces between fibers may be much more complex, including friction, crosslinks and several other physical processes that are challenging to simulate. Some of these may have major impact on the results, and others may not. This likely depends on the specific structure and loading conditions of the tissue. We carried out a friction sensitivity test in which we repeated the inverse fitting while using several friction coefficients. The results were very similar in stress-strain (less than 5% difference). We thus conclude that in the particular test reported herein ignoring friction did not adversely affect the accuracy of the model estimates substantially. Fiber-matrix interactions are potentially even more complex than fiber-fiber interactions given the wide diversity of components that form what we describe simply as “matrix”. These include non-fibrous components, such as GAG chains, that may act as lubricants and affect sliding, in which case their presence could make the tissue more compliant (Hatami-Marbini and Pachenari, 2020; Murienne et al., 2016). Other components include elastin fibers and cells. While we acknowledge the highly simplified fiber-fiber and fiber-matrix interactions we used in this study, we argue that the methodology we have described can be extended to incorporate much more complex interactions. Conventional continuum modeling techniques of the eye and other tissues simply ignore fiber-fiber and fiber-matrix interactions, yet it is not obvious to readers that these assumptions have been made, or how to relax them.

Second, we did not account for sub-fiber level features, such as collagen fiber crimp and various fiber or bundle diameters. Future work would benefit from introducing collagen fiber crimp (Grytz et al., 2014a) and various fiber diameters (Komai and Ushiki, 1991).

Third, our direct fiber model does not account for the very large number and complexity of fibers of the sclera. We posit that the fiber-level mechanics presented using the direct fiber model with reduced fiber density will likely be similar to the actual tissue. Therefore, our objective of introducing direct fiber model for simulating sclera is fulfilled. Meanwhile, it’s beneficial to use fewer fibers to reduce computational cost. We agree that in future work, it will be worthwhile to work on improving computational efficiency and building the model with more fibers.

Fourth, we did not account for regional variations. In our model structure we did not observe the strong well-aligned region of fibers in the radial direction that have been reported to take about 10% of the sclera nearest to the choroid. This could be due to variations between eyes, regional inhomogeneities (the radial fibers appear to be more readily distinguished closer to the optic nerve head) or due to the specific coronal sections used for the reconstruction. It is likely that other regions and other eyes will produce different models that respond differently to mechanical loading. We posit that our goal in this manuscript was modest, aiming to demonstrate it is possible to build a fiber-based model that produces a macroscopic mechanical behavior matching experimental tests. It will be beneficial to consider more features and variations in future work.

Fifth, the model used to demonstrate the direct fiber method was reconstructed from PLM and experimental data from different species. We posit that the impact was minor because our goal was to demonstrate the reconstruction and modeling methodology. It should be clear to readers that the process could be redone with all data from the same species. In this sense the key result from this work still stands: the direct fiber modeling technique can work. If the goal is to obtain fiber properties that are as accurate as possible for a given species, then using all consistent data is probably necessary. More samples will also be needed to account for variability.

Sixth, the matrix mechanical properties were kept constant at literature values and not optimized iteratively like the fiber properties. This was for the sake of simplicity. Although the matrix properties could impact the load bearing of the fibers and potentially influence the fiber material parameters fit, we believe the impact is minor. It is because the matrix is much softer than the fibers (Coudrillier et al., 2015b) and, consequently, the fibers are the principal load-bearing component (Girard et al., 2009a; Huang et al., 2013). The final results in our model show that the total reaction forces borne by the matrix are 5%-10% of those borne by the fibers at 2% of strain (Supplementary Figure 1). Therefore, we reasoned that the properties of the fibers will predominantly influence how the model behaves and the matrix stiffness has only a slight influence on the fiber material parameters derived. Nevertheless, it would be beneficial to consider matrix properties variations in future studies. Introducing matrix properties changes to the model may be necessary to properly account for the effects of age and interweaving, given the evidence that changes in matrix properties could result in age-related changes in sclera properties (Grytz et al., 2014b).

Seventh, although our model has similar boundary conditions to the experiment it is impossible to match the experiment precisely. Holding the tissues with clamps, rakes or hooks alters the boundary conditions, in ways that are extremely complicated to replicate computationally. Moreover, the experiments to which we fit the model were biaxial stretch tests. This is not the physiologic mechanical condition of sclera. The mechanical behavior of sclera under inflation or more complex modes could be different.

Eighth, we have chosen only one set of experimental data for estimating the material properties of the fibers. Given the complexity of the mechanical behavior of sclera, fitting only a biaxial stretch test does not guarantee that the model will generalize to other conditions. It may be necessary to fit simultaneously to several experimental tests, as has been done for continuum models (Thomas et al., 2019).

Ninth, the thickness of the direct fiber model was 0.35mm, but the sclera thickness in Eilaghi et al.’s experimental study was roughly 0.5mm. Although there is a discrepancy in thickness, we believe it will not affect the derived fiber material properties because these were obtained by fitting stress-strain curves.

## 5. Conclusion

We have shown the possibility of developing specimen-specific direct fiber model of sclera that can represent the sclera fibrous microstructure better than the previous continuum modeling approaches and allow accurate capture of sclera mechanics. We successfully built the model with long, continuous, interwoven fibers that takes into account the effects of fiber interweaving and fiber-fiber interactions. Our results have demonstrated that the direct fiber model can match the fiber orientations measured in high-resolution PLM images simultaneously in coronal and sagittal planes. The model properties can be optimized through inverse fitting to match experimental stress-strain responses. The estimated fiber elastic modulus is in good agreement with the literature. The direct fiber modeling methodology potentially has broad application to simulate other fiber-based tissues. Overall, the direct fiber modeling technique in this study is important for characterizing sclera collagen architecture at the fiber level, analyzing microstructural responses to macroscale mechanical loadings, and for understanding the scleral biomechanical environment.

## Reference

Behkam, R., Kollech, H.G., Jana, A., Hill, A., Danford, F., Howerton, S., Ram, S., Rodríguez, J.J., Utzinger, U., Girkin, C.A., Vande Geest, J.P., 2019. Racioethnic differences in the biomechanical response of the lamina cribrosa. Acta biomaterialia 88, 131–140.

Boote, C., Sigal, I.A., Grytz, R., Hua, Y., Nguyen, T.D., Girard, M.J.A., 2020. Scleral structure and biomechanics. Progress in retinal and eye research 74, 100773.

Brown, D.J., Morishige, N., Neekhra, A., Minckler, D.S., Jester, J.V., 2007. Application of second harmonic imaging microscopy to assess structural changes in optic nerve head structure ex vivo. Journal of biomedical optics 12, 024029.

Campbell, I.C., Coudrillier, B., Mensah, J., Abel, R.L., Ethier, C.R., 2015. Automated segmentation of the lamina cribrosa using Frangi’s filter: A novel approach for rapid identification of tissue volume fraction and beam orientation in a trabeculated structure in the eye. Journal of The Royal Society Interface 12, 20141009.

Coudrillier, B., Boote, C., Quigley, H.A., Nguyen, T.D., 2013. Scleral anisotropy and its effects on the mechanical response of the optic nerve head. Biomechanics and Modeling in Mechanobiology 12, 941–963.

Coudrillier, B., Geraldes, D.M., Vo, N.T., Atwood, R., Reinhard, C., Campbell, I.C., Raji, Y., Albon, J., Abel, R.L., Ethier, C.R., 2016. Phase-Contrast Micro-Computed Tomography Measurements of the Intraocular Pressure-Induced Deformation of the Porcine Lamina Cribrosa. IEEE transactions on medical imaging 35, 988–999.

Coudrillier, B., Pijanka, J.K., Jefferys, J.L., Goel, A., Quigley, H.A., Boote, C., Nguyen, T.D., 2015a. Glaucoma-related changes in the mechanical properties and collagen micro- architecture of the human sclera. PLoS One 10, e0131396.

Coudrillier, B., Pijanka, J.K., Jefferys, J.L., Sorensen, T., Quigley, H.A., Boote, C., Nguyen, T.D., 2015b. Collagen structure and mechanical properties of the human sclera: analysis for the effects of age. Journal of biomechanical engineering 137, 041006.

Coudrillier, B., Tian, J., Alexander, S., Myers, K.M., Quigley, H.A., Nguyen, T.D., 2012. Biomechanics of the human posterior sclera: age-and glaucoma-related changes measured using inflation testing. Investigative Ophthalmology & Visual Science 53, 1714–1728.

Danford, F.L., Yan, D., Dreier, R.A., Cahir, T.M., Girkin, C.A., Vande Geest, J.P., 2013. Differences in the region-and depth-dependent microstructural organization in normal versus glaucomatous human posterior sclerae. Investigative Ophthalmology & Visual Science 54, 7922–7932.

Eilaghi, A., Flanagan, J.G., Tertinegg, I., Simmons, C.A., Brodland, G.W., Ethier, C.R., 2010. Biaxial mechanical testing of human sclera. Journal of Biomechanics 43, 1696–1701.

Elliott, D.M., Setton, L.A., 2001. Anisotropic and inhomogeneous tensile behavior of the human anulus fibrosus: experimental measurement and material model predictions. Journal of biomechanical engineering 123, 256–263.

Ethier, C.R., Johnson, M., Ruberti, J., 2004. Ocular biomechanics and biotransport. Annu. Rev. Biomed. Eng. 6, 249–273.

Girard, M.J.A., Downs, J.C., Bottlang, M., Burgoyne, C.F., Suh, J.-K.F., 2009a. Peripapillary and Posterior Scleral Mechanics—Part II: Experimental and Inverse Finite Element Characterization. Journal of Biomechanical Engineering 131.

Girard, M.J.A., Downs, J.C., Burgoyne, C.F., Suh, J.-K.F., 2009b. Peripapillary and posterior scleral mechanics—part I: development of an anisotropic hyperelastic constitutive model. Journal of biomechanical engineering 131.

Girard, M.J.A., Suh, J.-K.F., Bottlang, M., Burgoyne, C.F., Downs, J.C., 2009c. Scleral Biomechanics in the Aging Monkey Eye. Investigative Ophthalmology & Visual Science 50, 5226–5237.

Gogola, A., Jan, N.-J., Brazile, B.L., Lam, P., Lathrop, K.L., Chan, K.C., Sigal, I.A., 2018a. Spatial patterns and age-related changes of the collagen crimp in the human cornea and sclera. Investigative Ophthalmology & Visual Science 59, 2987–2998.

Gogola, A., Jan, N.-J., Lathrop, K.L., Sigal, I.A., 2018b. Radial and circumferential collagen fibers are a feature of the peripapillary sclera of human, monkey, pig, cow, goat, and sheep. Investigative ophthalmology & visual science 59, 4763–4774.

Grytz, R., Fazio, M.A., Girard, M.J.A., Libertiaux, V., Bruno, L., Gardiner, S., Girkin, C.A., Downs, J.C., 2014a. Material properties of the posterior human sclera. Journal of the Mechanical Behavior of Biomedical Materials 29, 602–617.

Grytz, R., Fazio, M.A., Libertiaux, V., Bruno, L., Gardiner, S., Girkin, C.A., Downs, J.C., 2014b. Age-and race-related differences in human scleral material properties. Investigative Ophthalmology & Visual Science 55, 8163–8172.

Grytz, R., Krishnan, K., Whitley, R., Libertiaux, V., Sigal, I.A., Girkin, C.A., Downs, J.C., 2020a. A Mesh-Free Approach to Incorporate Complex Anisotropic and Heterogeneous Material Properties into Eye-Specific Finite Element Models. Comput Methods Appl Mech Eng 358.

Grytz, R., Meschke, G., 2009. Constitutive modeling of crimped collagen fibrils in soft tissues. Journal of the Mechanical Behavior of Biomedical Materials 2, 522–533.

Grytz, R., Meschke, G., 2010. A computational remodeling approach to predict the physiological architecture of the collagen fibril network in corneo-scleral shells. Biomech Model Mechanobiol 9, 225–235.

Grytz, R., Meschke, G., Jonas, J.B., 2011. The collagen fibril architecture in the lamina cribrosa and peripapillary sclera predicted by a computational remodeling approach. Biomechanics and modeling in mechanobiology 10, 371–382.

Grytz, R., Yang, H., Hua, Y., Samuels, B.C., Sigal, I.A., 2020b. Connective tissue remodeling in myopia and its potential role in increasing risk of glaucoma. Current Opinion in Biomedical Engineering 15, 40–50.

Guerin, H.L., Elliott, D.M., 2007. Quantifying the contributions of structure to annulus fibrosus mechanical function using a nonlinear, anisotropic, hyperelastic model. Journal of Orthopaedic Research 25, 508–516.

Hadi, M.F., Barocas, V.H., 2013. Microscale Fiber Network Alignment Affects Macroscale Failure Behavior in Simulated Collagen Tissue Analogs. Journal of Biomechanical Engineering 135.

Hatami-Marbini, H., Pachenari, M., 2020. The contribution of sGAGs to stress-controlled tensile response of posterior porcine sclera. PLoS One 15, e0227856.

Hoerig, C., McFadden, S., Hoang, Q.V., Mamou, J., 2022. Biomechanical changes in myopic sclera correlate with underlying changes in microstructure. Experimental Eye Research, 109165.

Holzapfel, G.A., 2001. Biomechanics of soft tissue. The handbook of materials behavior models 3, 1049–1063.

Hua, Y., Voorhees, A.P., Jan, N.-J., Wang, B., Waxman, S., Schuman, J.S., Sigal, I.A., 2020. Role of radially aligned scleral collagen fibers in optic nerve head biomechanics. Experimental Eye Research 199, 108188.

Huang, W., Fan, Q., Wang, W., Zhou, M., Laties, A.M., Zhang, X., 2013. Collagen: a potential factor involved in the pathogenesis of glaucoma. Med Sci Monit Basic Res 19, 237–240.

Huang, X., Zhou, Q., Liu, J., Zhao, Y., Zhou, W., Deng, D., 2017. 3D stochastic modeling, simulation and analysis of effective thermal conductivity in fibrous media. Powder technology 320, 397–404.

Islam, M.R., Picu, R.C., 2018. Effect of Network Architecture on the Mechanical Behavior of Random Fiber Networks. Journal of Applied Mechanics 85.

Jan, N.-J., Brazile, B.L., Hu, D., Grube, G., Wallace, J., Gogola, A., Sigal, I.A., 2018. Crimp around the globe; patterns of collagen crimp across the corneoscleral shell. Experimental Eye Research 172, 159–170.

Jan, N.-J., Gomez, C., Moed, S., Voorhees, A.P., Schuman, J.S., Bilonick, R.A., Sigal, I.A., 2017a. Microstructural crimp of the lamina cribrosa and peripapillary sclera collagen fibers. Investigative Ophthalmology & Visual Science 58, 3378–3388.

Jan, N.-J., Grimm, J.L., Tran, H., Lathrop, K.L., Wollstein, G., Bilonick, R.A., Ishikawa, H., Kagemann, L., Schuman, J.S., Sigal, I.A., 2015. Polarization microscopy for characterizing fiber orientation of ocular tissues. Biomed. Opt. Express 6, 4705–4718.

Jan, N.-J., Lathrop, K.L., Sigal, I.A., 2017b. Collagen architecture of the posterior pole: high-resolution wide field of view visualization and analysis using polarized light microscopy. Investigative ophthalmology & visual science 58, 735–744.

Jan, N.-J., Lee, P.-Y., Wallace, J., Iasella, M., Gogola, A., Sigal, I.A., 2022. Stretch-Induced Uncrimping of Equatorial Sclera Collagen Bundles. bioRxiv, 2022.2009.2013.507860.

Jan, N.-J., Sigal, I.A., 2018. Collagen fiber recruitment: A microstructural basis for the nonlinear response of the posterior pole of the eye to increases in intraocular pressure. Acta Biomaterialia 72, 295–305.

Kollech, H.G., Ayyalasomayajula, A., Behkam, R., Tamimi, E., Furdella, K., Drewry, M., Vande Geest, J.P., 2019. A Subdomain Method for Mapping the Heterogeneous Mechanical Properties of the Human Posterior Sclera. Frontiers in Bioengineering and Biotechnology 7.

Komai, Y., Ushiki, T., 1991. The three-dimensional organization of collagen fibrils in the human cornea and sclera. Investigative Ophthalmology & Visual Science 32, 2244–2258.

Lagarias, J.C., Reeds, J.A., Wright, M.H., Wright, P.E., 1998. Convergence properties of the Nelder--Mead simplex method in low dimensions. SIAM Journal on optimization 9, 112–147.

Lanir, Y., 2017. Multi-scale Structural Modeling of Soft Tissues Mechanics and Mechanobiology. Journal of Elasticity 129, 7–48.

Lee, P.-Y., Yang, B., Hua, Y., Waxman, S., Zhu, Z., Ji, F., Sigal, I.A., 2022a. Real-time imaging of optic nerve head collagen microstructure and biomechanics using instant polarized light microscopy. Experimental Eye Research 217, 108967.

Lee, P.-Y., Yang, B., Sigal, I.A., 2022b. Quantitative stretch-induced collagen fiber recruitment and microarchitecture changes using instant polarized light microscopy, 15th World Congress on Computational Mechanics, Hybrid conference, presentation delivered remotely, in-person held in Yokohama Japan, July 31-August 5, 2022.

Licup, A.J., Münster, S., Sharma, A., Sheinman, M., Jawerth, L.M., Fabry, B., Weitz, D.A., MacKintosh, F.C., 2015. Stress controls the mechanics of collagen networks. Proceedings of the National Academy of Sciences 112, 9573–9578.

Ling, Y.T.T., Shi, R., Midgett, D.E., Jefferys, J.L., Quigley, H.A., Nguyen, T.D., 2019. Characterizing the collagen network structure and pressure-induced strains of the human lamina cribrosa. Investigative Ophthalmology & Visual Science 60, 2406–2422.

Matuschke, F., Amunts, K., Axer, M., 2021. fastPLI: A Fiber Architecture Simulation Toolbox for 3D-PLI. Journal of Open Source Software 6, 3042.

Matuschke, F., Ginsburger, K., Poupon, C., Amunts, K., Axer, M., 2019. Dense fiber modeling for 3D-Polarized Light Imaging simulations. Advances in parallel computing 34, 240–253.

Miles, J., 2005. R-squared, adjusted R-squared. Encyclopedia of statistics in behavioral science.

Moerman, K.M., 2018. GIBBON: the geometry and image-based bioengineering add-on. Journal of Open Source Software 3, 506.

Murienne, B.J., Chen, M.L., Quigley, H.A., Nguyen, T.D., 2016. The contribution of glycosaminoglycans to the mechanical behaviour of the posterior human sclera. Journal of The Royal Society Interface 13.

Nerurkar, N.L., Mauck, R.L., Elliott, D.M., 2011. Modeling interlamellar interactions in angle-ply biologic laminates for annulus fibrosus tissue engineering. Biomechanics and modeling in mechanobiology 10, 973–984.

Petsche, S.J., Pinsky, P.M., 2013. The role of 3-D collagen organization in stromal elasticity: a model based on X-ray diffraction data and second harmonic-generated images. Biomechanics and modeling in mechanobiology 12, 1101–1113.

Picu, R.C., Deogekar, S., Islam, M.R., 2018. Poisson’s Contraction and Fiber Kinematics in Tissue: Insight From Collagen Network Simulations. Journal of Biomechanical Engineering 140.

Pijanka, J.K., Coudrillier, B., Ziegler, K., Sorensen, T., Meek, K.M., Nguyen, T.D., Quigley, H.A., Boote, C., 2012. Quantitative Mapping of Collagen Fiber Orientation in Non-glaucoma and Glaucoma Posterior Human Sclerae. Investigative Ophthalmology & Visual Science 53, 5258–5270.

Pijanka, J.K., Markov, P.P., Midgett, D., Paterson, N.G., White, N., Blain, E.J., Nguyen, T.D., Quigley, H.A., Boote, C., 2019. Quantification of collagen fiber structure using second harmonic generation imaging and two-dimensional discrete Fourier transform analysis: Application to the human optic nerve head. Journal of biophotonics 12, e201800376.

Pijanka, J.K., Spang, M.T., Sorensen, T., Liu, J., Nguyen, T.D., Quigley, H.A., Boote, C., 2015. Depth-dependent changes in collagen organization in the human peripapillary sclera. PloS one 10, e0118648.

Roberts, M.D., Liang, Y., Sigal, I.A., Grimm, J.L., Reynaud, J., Bellezza, A., Burgoyne, C.F., Downs, J.C., 2010. Correlation between local stress and strain and lamina cribrosa connective tissue volume fraction in normal monkey eyes. Investigative ophthalmology & visual science 51, 295–307.

Saiman, M., Wahab, M., Wahit, M., 2014. The effect of fabric weave on the tensile strength of woven kenaf reinforced unsaturated polyester composite, Proceedings of the International Colloquium in Textile Engineering, Fashion, Apparel and Design 2014 (ICTEFAD 2014). Springer, pp. 25–29.

Schwaner, S.A., Hannon, B.G., Feola, A.J., Ethier, C.R., 2020a. Biomechanical properties of the rat sclera obtained with inverse finite element modeling. Biomechanics and Modeling in Mechanobiology 19, 2195–2212.

Schwaner, S.A., Perry, R.N., Kight, A.M., Winder, E., Yang, H., Morrison, J.C., Burgoyne, C.F., Ross Ethier, C., 2020b. Individual-Specific Modeling of Rat Optic Nerve Head Biomechanics in Glaucoma. Journal of Biomechanical Engineering 143.

Sigal, I.A., Flanagan, J.G., Tertinegg, I., Ethier, C.R., 2004. Finite Element Modeling of Optic Nerve Head Biomechanics. Investigative Ophthalmology & Visual Science 45, 4378–4387.

Sigal, I.A., Grimm, J.L., Jan, N.-J., Reid, K., Minckler, D.S., Brown, D.J., 2014. Eye-specific IOP-induced displacements and deformations of human lamina cribrosa. Investigative Ophthalmology & Visual Science 55, 1–15.

Stig, F., Hallström, S., 2019. Effects of crimp and textile architecture on the stiffness and strength of composites with 3D reinforcement. Advances in Materials Science and Engineering 2019.

Summers Rada, J.A., Shelton, S., Norton, T.T., 2006. The sclera and myopia. Experimental Eye Research 82, 185–200.

Thomas, V.S., Lai, V., Amini, R., 2019. A computational multi-scale approach to investigate mechanically-induced changes in tricuspid valve anterior leaflet microstructure. Acta Biomaterialia 94, 524–535.

Van Der Rijt, J.A., Van Der Werf, K.O., Bennink, M.L., Dijkstra, P.J., Feijen, J., 2006. Micromechanical testing of individual collagen fibrils. Macromolecular bioscience 6, 697–702.

van Tonder, J.D., Venter, M., Venter, G., 2023. A Novel Method for Resolving Non-Unique Solutions Observed in Fitting Parameters to the Mooney Rivlin Material Model. Available at SSRN 4364489.

Voorhees, A.P., Jan, N.-J., Hua, Y., Yang, B., Sigal, I.A., 2018. Peripapillary sclera architecture revisited: a tangential fiber model and its biomechanical implications. Acta Biomaterialia 79, 113–122.

Voorhees, A.P., Jan, N.-J., Sigal, I.A., 2017. Effects of collagen microstructure and material properties on the deformation of the neural tissues of the lamina cribrosa. Acta biomaterialia 58, 278–290.

Wagner, D.R., Lotz, J.C., 2004. Theoretical model and experimental results for the nonlinear elastic behavior of human annulus fibrosus. Journal of orthopaedic research 22, 901–909.

Wang, B., Hua, Y., Brazile, B.L., Yang, B., Sigal, I.A., 2020. Collagen fiber interweaving is central to sclera stiffness. Acta Biomaterialia 113, 429–437.

Wenger, M.P.E., Bozec, L., Horton, M.A., Mesquida, P., 2007. Mechanical Properties of Collagen Fibrils. Biophysical Journal 93, 1255–1263.

Winkler, M., Jester, B., Nien-Shy, C., Massei, S., Minckler, D.S., Jester, J.V., Brown, D.J., 2010. High resolution three-dimensional reconstruction of the collagenous matrix of the human optic nerve head. Brain research bulletin 81, 339–348.

Yang, B., Brazile, B.L., Jan, N.-J., Hua, Y., Wei, J., Sigal, I.A., 2018a. Structured polarized light microscopy for collagen fiber structure and orientation quantification in thick ocular tissues. Journal of biomedical optics 23, 1–10.

Yang, B., Jan, N.-J., Brazile, B.L., Voorhees, A.P., Lathrop, K.L., Sigal, I.A., 2018b. Polarized light microscopy for 3-dimensional mapping of collagen fiber architecture in ocular tissues. Journal of biophotonics 11, e201700356.

Yang, B., Lee, P.-Y., Hua, Y., Brazile, B.L., Waxman, S., Ji, F., Zhu, Z., Sigal, I.A., 2021. Instant polarized light microscopy for imaging collagen microarchitecture and dynamics. Journal of biophotonics 14, e202000326.

Yang, L., Van Der Werf, K.O., Koopman, B.F., Subramaniam, V., Bennink, M.L., Dijkstra, P.J., Feijen, J., 2007. Micromechanical bending of single collagen fibrils using atomic force microscopy. Journal of Biomedical Materials Research Part A 82, 160–168.

Zhang, L., Albon, J., Jones, H., Gouget, C.L., Ethier, C.R., Goh, J.C., Girard, M.J., 2015. Collagen microstructural factors influencing optic nerve head biomechanics. Investigative Ophthalmology & Visual Science 56, 2031–2042.

Zhang, L., Lake, S.P., Lai, V.K., Picu, C.R., Barocas, V.H., Shephard, M.S., 2013. A coupled fiber-matrix model demonstrates highly inhomogeneous microstructural interactions in soft tissues under tensile load. Journal of biomechanical engineering 135, 011008.

Zhou, D., Abass, A., Eliasy, A., Studer, H.P., Movchan, A., Movchan, N., Elsheikh, A., 2019. Microstructure-based numerical simulation of the mechanical behaviour of ocular tissue. Journal of the Royal Society Interface 16, 20180685.

