## Supplementary Table for "A direct fiber approach to model sclera collagen architecture and biomechanics"

**Supplementary Table 1**

| Set # | C_10_ (MPa) | C_01_ (MPa) | Shear modulus (MPa) |
| --- | --- | --- | --- |
| 1 | 5732.2650 | -4989.8000 | 1484.93 |
| 2 | 5929.2900 | -5169.5800 | 1519.42 |
| 3 | 5971.4700 | -5230.2900 | 1482.36 |
| 4 | 5722.4110 | -4981.2290 | 1482.36 |
| 5 | 881.8312 | -100.0000 | 1563.66 |
| 6 | 119.4267 | 662.3430 | 1563.54 |
| 7 | 119.3580 | 662.0210 | 1562.76 |
| 8 | 5746.9000 | -5002.6000 | 1488.60 |
| 9 | 119.4156 | 662.3429 | 1563.52 |
| 10 | 7881.8000 | -7140.0000 | 1483.60 |
| 11 | 118.9000 | 659.5000 | 1556.80 |
| Max | 7881.8000 | 662.3430 | 1563.66 |
| Min | 118.9000 | -7140.0000 | 1482.36 |
| Average | 3485.7334 | -2724.2993 | 1522.87 |
| STDEV | 3142.4550 | 3160.9742 | 38.94 |

**Table 1.** 11 sets of C_10_ and C_01_ hyperelastic Mooney Rivlin material parameters that led to stress-strain curves in good agreement with the experimental data from the literature (see main text). The resultant fiber shear modulus is with an average of 1522.87 MPa and a standard deviation of 38.94 MPa.
