## Supplementary Figure for "A direct fiber approach to model sclera collagen architecture and biomechanics"

**Supplementary Figure 1**

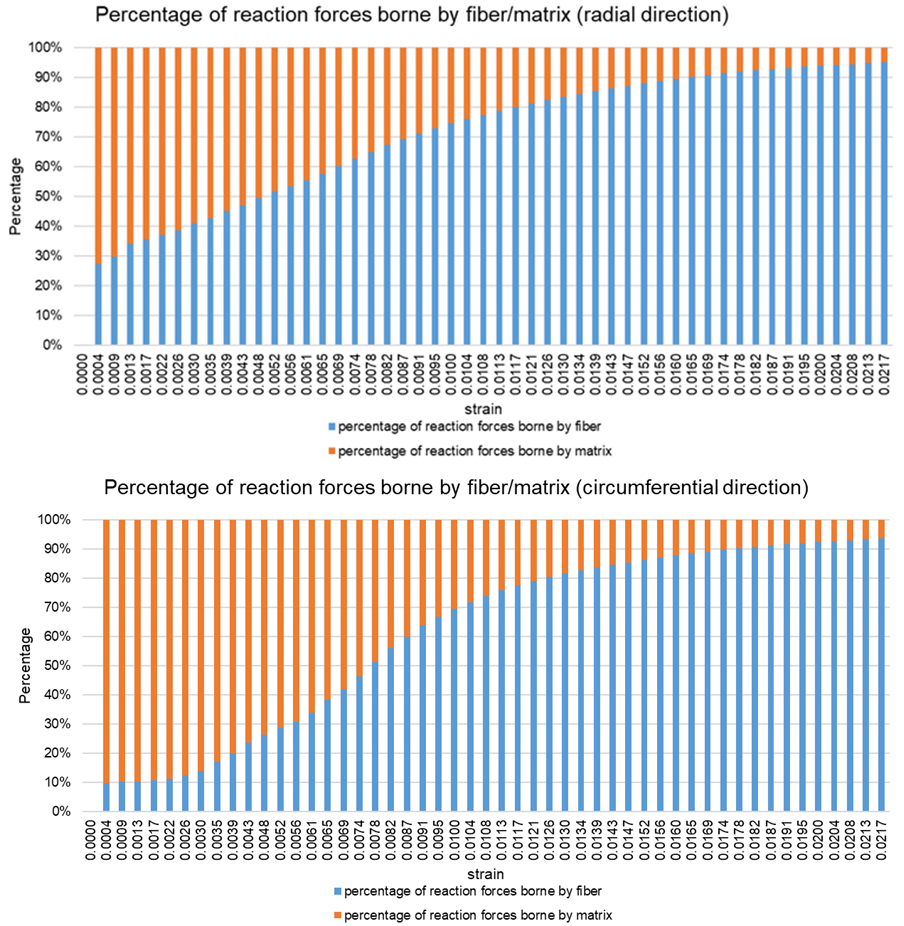

**Figure 1.** Total reaction forces borne by the matrix and fibers, respectively.
